# Hypoxia Dynamically Regulates DBC1 Ubiquitination and Stability by SIAH2 and OTUD5 in Breast Cancer Progression

**DOI:** 10.1101/2022.01.11.475808

**Authors:** Qiangqiang Liu, Qian Luo, Jianyu Feng, Yanping Zhao, Linlin Liu, Biao Ma, Hongcheng Cheng, Tian Zhao, Jiaojiao Zhang, Chenglong Mu, Linbo Chen, Hong Lei, Yijia Long, Jingyi Su, Guo Chen, Yanjun Li, Quan Chen, Yushan Zhu

**Affiliations:** State Key Laboratory of Medicinal Chemical Biology, Frontiers Science Center for Cell Responses, Tianjin Key Laboratory of Protein Science, College of Life Sciences, Nankai University, Tianjin, China; School of Statistics and Data Science, LPMC and KLMDASR, Nankai University, Tianjin, China

**Author notes:** Corresponding author: Yushan Zhu.

## Abstract

DBC1 has been characterized as a key regulator of physiological and pathophysiological activities, such as DNA damage, senescence and tumorigenesis. However, the mechanism by which the functional stability of DBC1 is regulated has yet to be elucidated. Here, we report that the ubiquitination-mediated degradation of DBC1 is dynamically regulated by the E3 ubiquitin ligase SIAH2 and deubiquitinase OTUD5 under hypoxic stress. Mechanistically, hypoxia promoted the competitive binding of SIAH2 with OTUD5 to DBC1, resulting in the ubiquitination and subsequent degradation of DBC1 through the ubiquitin–proteasome pathway. *Siah2* knockout inhibited tumor cell proliferation and migration, which could be rescued by double knockout of *Siah2/DBC1*. Human tissue microarray analysis further revealed that the SIAH2/DBC1 axis was responsible for tumor progression under hypoxic stress. These findings define a key role of the hypoxia-mediated SIAH2-DBC1 pathway in the progression of human breast cancer and provide novel insights into the metastatic mechanism of breast cancer.

## Introduction

The occurrence and development of tumors is modulated by the dual regulation of genetic instability and the tumor microenvironment (Singleton, Macann, and Wilson 2021). Hypoxic stress or low oxygen tension, a major hallmark of the tumor microenvironment, plays an essential role in the progression and metastasis of many solid tumors (Cheng et al. 2020; Lee et al. 2019). Moreover, our previous studies have documented that hypoxic stress attenuates tumorigenesis and progression by modulating the Hippo signaling pathway and mitochondrial biogenesis (Ma et al. 2015; Ma et al. 2019). To extensively understand the critical roles of hypoxic stress in tumorigenesis or tumor development, more precise mechanisms need to be further explored.

Deleted in Breast Cancer 1 (DBC1; also known as CCAR2) is a nuclear protein containing multifunctional domains and plays a critical role in a variety of cancers. Importantly, DBC1 cooperates with lots of epigenetic and transcriptional factors to regulate cell activities. DBC1 mediates p53 function not only by inhibiting the deacetylase activity of SIRT1, but also by inhibition of p53 ubiquitination and degradation (Qin et al. 2015; Akande et al. 2019) (Kim, Chen, and Lou 2008; Zhao et al. 2008). In addition, DBC1 can specifically inhibit deacetylase HDAC3 activity and alter its subcellular distribution, resulting in cell senescence (Chini et al. 2010). Most importantly, mice with a genetic deletion of *DBC1* exhibited more likely to spontaneously develop lung tumors, liver tumors, lymphomas, and teratomas and showed poor overall survival (Qin et al. 2015). Recently, the downregulation of DBC1 highly correlates with poor prognosis and distant metastatic relapse in breast, colon, and prostate cancer patients, and low levels of DBC1 determine tumor grade and metastasis (Won et al. 2015; Noguchi et al. 2014; Yu et al. 2013). Accumulating evidence has shown that DBC1 activity and quality of control play key roles in tumorigenicity, while the mechanisms by which DBC1 stability is regulated remain elusive.

Protein ubiquitination is one of the most-studied posttranslational modifications and is critical and essential for protein stability, activity, localization and biological function. Here, we delineated that the ubiquitination and stability of DBC1 were dynamically orchestrated by the E3 ubiquitin ligase SIAH2 and deubiquitinase OTUD5 under hypoxic stress. Mechanistically, hypoxic stress promoted the interaction between DBC1 and SIAH2 and enhanced the disassociation of DBC1 from OTUD5, resulting in an increase in DBC1 ubiquitination and degradation, contributing to tumor cell proliferation and migration. Human tissue microarray analysis further revealed that the SIAH2/DBC1 axis was responsible for regulating tumor progression under hypoxic stress. Our findings provide novel insights into the metastatic mechanism of breast cancer and a promising therapeutic target for breast cancer.

## Results

### Hypoxic stress triggers the degradation of DBC1

Hypoxia is a common hallmark of solid tumors and contributes to the development and progression of many cancers (Lee et al. 2019). To investigate the effects of hypoxia on breast cancer cells, we first performed RNA-seq analysis of MDA-MB-231 cells in response to hypoxic stress. Differential expression analysis showed that 1151 genes were significantly upregulated and 310 genes were downregulated (q < 0.05) under hypoxic stress (Figure 1A). Enrichment analysis of differentially expressed genes by Metascape suggested that the upregulated genes were related to the pathways of cell proliferation and cancer; in contrast, the downregulated genes were implicated in the DNA damage repair, senescence, SIRT1 and p53 signaling pathways (Figure 1B). To further investigate the mechanism by which hypoxia regulates the p53 signaling pathway, we examined p53 pathway activity by western blotting, and found that the stabilities of SIRT1 and p53 protein were not changed under hypoxia, but the acetylation of p53 was decreased (Figure 1C). Interestingly, we observed that the protein level of DBC1 gradually decreased with the duration of hypoxia (Figure 1D), which could be inhibited by the proteasome inhibitor MG132 (Figure 1E), suggesting that hypoxia induced DBC1 degradation through the ubiquitin-proteasome system. In most cell lines derived from different sub-types of breast cancer or other cancers, DBC1 protein was degraded under hypoxia, suggesting the reduced stability of DBC1 was a general rule for tumor cell survival in hypoxic environment (Figure 1F and Supplementary file 1), while the mRNA level of DBC1 was not changed under hypoxia (Figure 1G and Supplementary file 1). Similarly, the RNA-Seq data sets of *DBC1* knockout MDA-MB-231 cells showed that genes related to cell proliferation and migration were also upregulated, and genes associated with DNA repair and apoptosis were downregulated (Figure 1H, I and Figure 1-figure supplement 1A, B). Collectively, these results suggest that hypoxia-induced DBC1 degradation contributes to tumor progression by regulating multiple pathways.

**Figure 1.**
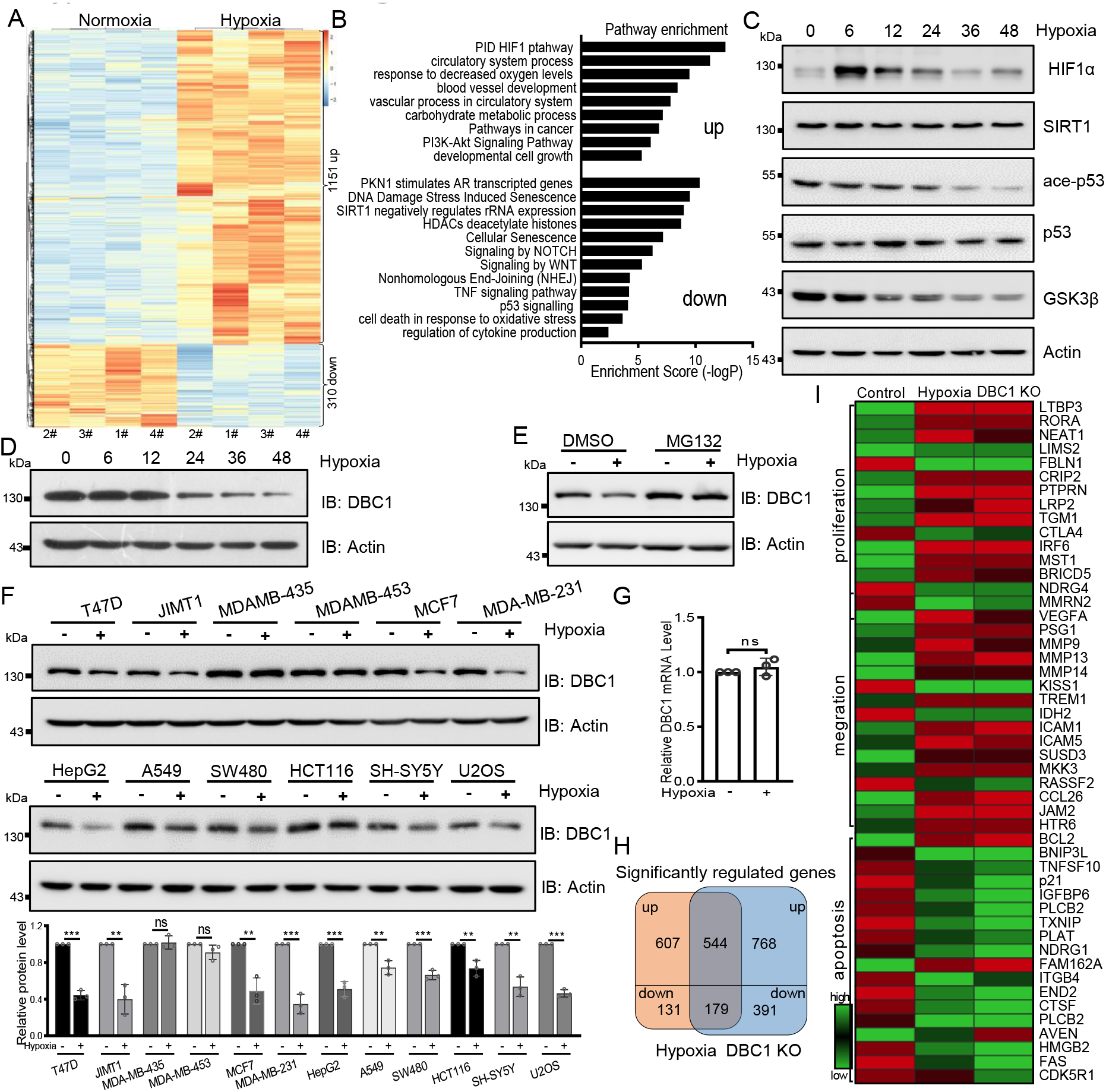
Hypoxia induces DBC1 degradation. **A,** Heatmap showing differential transcriptomic expression compared with normoxia and hypoxic stress in MDA-MB-231 cells. **B,** Highlights of the enriched pathways from transcriptomic expression by Metascape. **C,** Western blotting analysis of the protein levels using the indicated antibodies over time in MDA-MB-231 cells under hypoxic stress. **D,** Protein levels of DBC1 were detected by western blotting. **E,** Hypoxia-mediated DBC1 degradation is inhibited by the proteasomal inhibitor MG132 (10 μM). **F,** Western blotting analysis of the degradation of DBC1 in the indicated cell lines induced by hypoxia. mean ± SEM from three independent experiments, Student’s t test, *P < 0.05, **P < 0.01. **G,** Relative mRNA levels of DBC1 in MDA-MB-231 cells under hypoxia or normoxia were analysed using qRT–PCR. Data are shown as the mean ± SEM. Statistical significance was calculated using two-way ANOVA. **H,** Venn diagram showing the numbers of DBC1-regulated genes common to hypoxia target genes. **I,** Heatmap showing the representatives of differentially regulated transcripts associated with tumor initiation and progression.

### SIAH2 interacts with DBC1 and regulates its stability

To identify the E3 ligases that potentially ubiquitinates DBC1 to contribute its degradation, several ubiquitin E3 ligases previously reported to be involved in the hypoxia response were screened, and SIAH2 was finally identified to be responsible for the stability of DBC1 (Figure 2A). To confirm our hypothesis, LC–MS/MS analysis of proteins bound to SIAH2^RM^ (a SIAH2 enzyme inactive mutant) was performed. DBC1 was validated as one of the substrates of SIAH2 (Figure 2B). To further explore the relationship between SIAH2 and DBC1, we carried out a Co-IP assay and found that exogenously expressed Flag-SIAH2^RM^ and Myc-DBC1 were reciprocally coimmunoprecipitated (Figure 2-figure supplement 1A, B). Meanwhile, we validated the interaction between endogenous SIAH2 and DBC1 (Figure 2C), and purified glutathione S-transferase (GST) SIAH2 was able to pull down His-DBC1 *in vitro* (Figure 2D), suggesting that DBC1 could directly interact with SIAH2. To identify the necessary domain of DBC1 responsible for interaction with SIAH2, we constructed several truncations of DBC1 and SIAH2 (Figure 2-figure supplement 1C, D), and the Co-IP assay showed that the 1-230 amino acids at the N-terminus of DBC1 were necessary for binding to SIAH2 (Figure 2E), and the fulllength SIAH2 was required for their interaction (Figure 2F). Importantly, the N-terminus (1-461) of the DBC1 protein rather than the C-terminus (462-923) was pulled down by SIAH2 *in vitro* (Figure 2G, H). These results suggest that the N-terminus of DBC1 is crucial for its interaction with SIAH2. In line with the fact that SIAH2, as an E3 ligase, mediates substrate degradation by the ubiquitin–proteasome pathway, we found that SIAH2, rather than SIAH2^RM^, reduced the protein level of endogenous DBC1 in a dose-dependent manner (Figure 2I, J) and had no effect on the transcriptional level of DBC1 (Figure 2-figure supplement 1E and Supplementary file 2). A CHX chase assay showed that ectopic expression of wild-type SIAH2, but not SIAH2^RM^, promoted DBC1 degradation over time (Figure 2K, L and Supplementary file 2). Furthermore, the proteasome inhibitor MG132 completely reversed DBC1 destabilization (Figure 2M), whereas the lysosomal inhibitor bafilomycin A1 (BA1) had no such function (Figure 2-figure supplement 1F), suggesting that SIAH2 mediated DBC1 degradation through the proteasome pathway rather than lysosomes. These results reveal that SIAH2 promotes DBC1 degradation by directly interacting with DBC1.

**Figure 2.**
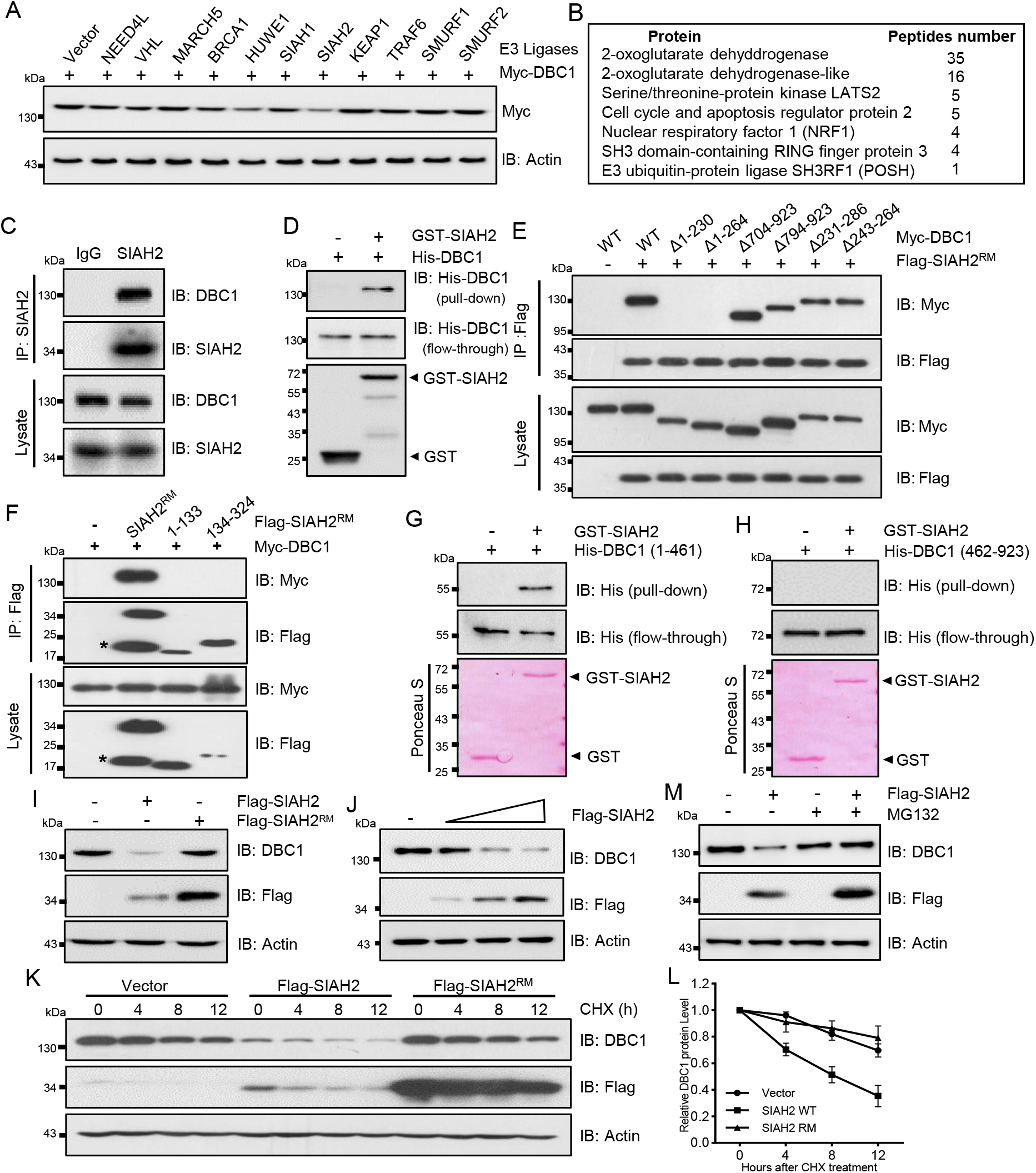
SIAH2 interacts with DBC1 and regulates its stability. **A,** Screening of the potential E3 ligase of DBC1 in HeLa cells cotransfected with Myc-DBC1 and the indicated plasmids into HeLa cells for 24 h, and Myc-DBC1 protein levels were detected by western blotting with an anti-Myc antibody. **B,** List of representative proteins identified by SIAH2 Co-IP/MS, and the number of peptides for each protein-identified peptide is indicated. **C,** Co-IP analysis of endogenous interaction between DBC1 and SIAH2 by the indicated antibodies. **D,** GST pulldown analysis showed the direct interactions between bacterially expressed GST-SIAH2 and His-DBC1 *in vitro*. **E,** Mapping the domain of DBC1 interacting with Flag-SIAH2^RM^ in HEK293T cells cotransfected with full-length or truncated forms of Myc-DBC1 and Flag-SIAH2^RM^. **F,**Mapping the domain of SIAH2 interacting with Myc-DBC1 in HEK293T cells that were cotransfected with full-length or truncated forms of Myc-DBC1 and Flag-SIAH2^RM^, * represents a nonspecific band. **G-H,** GST pulldown analysis showing the direct interaction between bacterially expressed GST-SIAH2 and the N- or C-terminal half of His-DBC1 *in vitro.* **I,** Western blotting analysis showing the protein level of endogenous DBC1 in HeLa cells exogenously expressing Flag-SIAH2 or Flag-SIAH2^RM^. **J-K,** Western blotting analysis showing the half-life of endogenous DBC1 in HeLa cells transfected with Flag-SIAH2 or Flag-SIAH2^RM^ for 24 h and treated with CHX (10 μM) at the indicated time (**J**). Quantification of the DBC1 protein level as shown (**K**). **L,** Western blotting analysis showing the DBC1 protein level in HeLa cells transfected with SIAH2 by quantitative gradient. **M,** Western blotting analysis showing the DBC1 level in HeLa cells overexpressing Flag-SIAH2 and treated with the proteasomal inhibitor MG132.

### SIAH2 is responsible for DBC1 ubiquitination and degradation under hypoxic stress

Next, we checked whether SIAH2 ubiquitinated DBC1, and the results showed that ectopic expression of SIAH2, but not SIAH2^RM^, dramatically increased the ubiquitination level of DBC1 (Figure 3A, B). Additionally, immunoprecipitation analysis proved that SIAH2 induced the polyubiquitination of DBC1 in the form of K48 conjugation (Figure 3-figure supplement 1A, B). *In vitro* ubiquitination assay further revealed that DBC1 was directly ubiquitinated by SIAH2 rather than the E3 ligase-dead mutant SIAH2^RM^ (Figure 3C). To identify the sites of DBC1 ubiquitinated by SIAH2, we performed MS analysis, and the results showed that K287 of DBC1 was the potential site ubiquitinated by SIAH2 (Figure 3D). Next, we found that ectopic expression of SIAH2 did not increase the ubiquitination level of the K287R mutant of DBC1, and the mutant did not degrade (Figure 3E, Figure 3-figure supplement 1C, D). In conclusion, these results demonstrate that SIAH2 ubiquitinates DBC1 to mediate its degradation. To further understand the mechanism underlying hypoxia-induced DBC1 degradation, we examined whether SIAH2 was responsible for DBC1 ubiquitination and degradation in response to hypoxia. Our results showed that under hypoxic stress, both *SIAH2* deletion and K287R mutation of DBC1 failed to increase DBC1 ubiquitination (Figure 3F, G), and DBC1 degradation was also inhibited (Figure 3H-J and Supplementary file 3). Consistently, knockout of *SIAH2* prolonged the half-life of DBC1 under hypoxia (Figure 3K, L and Supplementary file 3). Taken together, these results demonstrate that hypoxia-induced DBC1 degradation was dependent on SIAH2-mediated DBC1 ubiquitination.

**Figure 3.**
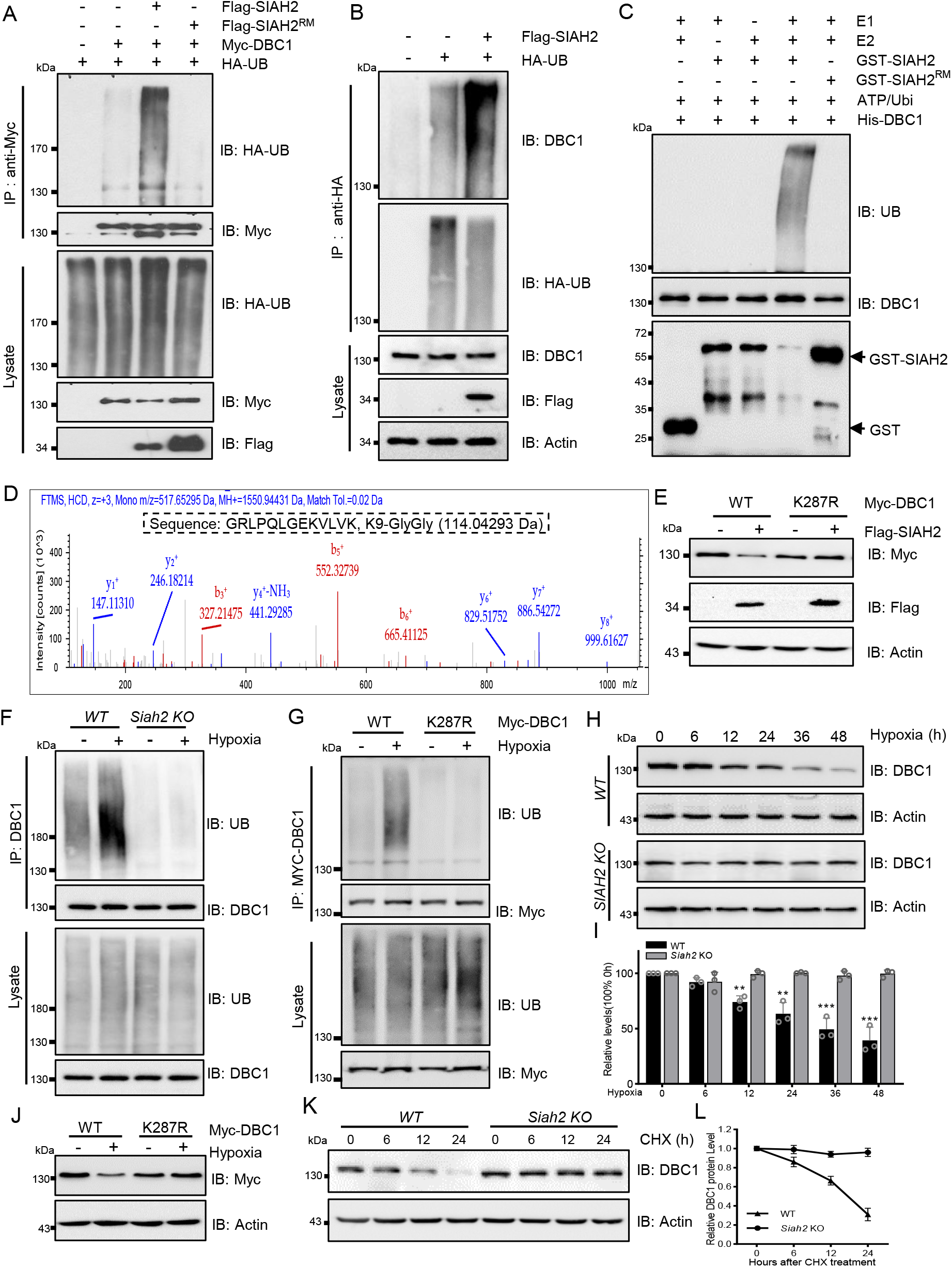
SIAH2 promotes DBC1 polyubiquitination and degradation under hypoxia. **A,** Co-IP analysis of DBC1 ubiquitination levels in HEK293T cells cotransfected with HA-Ub and wild-type SIAH2 or SIAH2^RM^. **B,** Co-IP analysis of ubiquitinated DBC1 in HEK293T cells cotransfected with HA-Ub and Flag-SIAH2. **C,** *In vitro* ubiquitination analysis of His-DBC1 coincubated with purified SIAH2 rather than SIAH2^RM^. **D,** LC–MS/MS analysis of the ubiquitinated sites in DBC1. **E,** Western blotting analysis showing the DBC1 level in HeLa cells cotransfected with Flag-SIAH2 and wild-type DBC1 or DBC1-K287R. **F,** Co-IP analysis showing the ubiquitination level of endogenous DBC1 in wild-type and *Siah2* knockout MDA-MB-231 cells under hypoxia. **G,** Co-IP analysis showing the ubiquitination level of exogenous DBC1 in HeLa cells transfected with wild-type DBC1 or DBC1-K287R under hypoxia. **H,** Western blotting analysis showing the endogenous DBC1 protein level in wild-type and *Siah2* knockout MDA-MB-231 cells under hypoxia. **I,** DBC1 protein levels from (**H**) were quantified (mean ± SEM; n=3 biologically independent samples; **P < 0.01, ***P < 0.001) **J,** Western blotting analysis showing the protein level of exogenous DBC1 in HeLa cells transfected with wild-type DBC1 or DBC1-K287R under hypoxia. **K-L,** Western blotting analysis showing the half-life of endogenous DBC1 in wild-type and *Siah2* knockout MDA-MB-231 cells treated with CHX under hypoxia (**J**). Quantification of DBC1 protein levels is shown (**L**).

### OTUD5 regulates the deubiquitination and stability of DBC1

Strikingly, we also observed that the ubiquitination level of DBC1 was sharply decreased when returned to normal conditions after hypoxic stress (Figure 4A). Therefore, we screened the deubiquitinating enzymes (DUBs) plasmids library to identify the candidate responsible for deubiquitination of DBC1 (Figure 4-figure supplement 1A). Lots of interaction assays suggested that the potential DUB OTUD5 might interact with DBC1. Co-IP analysis demonstrated the exogenous interaction between OTUD5 and DBC1 (Figure 4-figure supplement 1B, C). Furthermore, we confirmed that endogenous DBC1 could directly interact with OTUD5 (Figure 4B), which was further verified by an *in vitro* pull-down assay using purified His-tagged DBC1 to pull down OTUD5 from MDA-MB-231 cell lysates (Figure 4C). We next tested whether DBC1 deubiquitination was dependent on OTUD5 enzyme activity. The ubiquitination assay showed that only ectopic expression of wild-type OTUD5 but not inactive mutant C224S led to DBC1 ubiquitination level decrease under hypoxia (Figure 4D). Consistently, hypoxia-induced endogenous DBC1 degradation was specifically inhibited by expression of wild-type OTUD5 rather than the inactive mutant (Figure 4E). Moreover, the CHX assay also confirmed that ectopic expression of wild-type OTUD5 obviously prolonged the half-life of DBC1 (Figure 4F, G and Supplementary file 4), indicating that deubiquitination of DBC1 was beneficial for its stability. Domain mapping analysis further revealed that the N-terminal region (amino acid residues 1-230) of DBC1, which binds with SIAH2, was also necessary for its interaction with OTUD5 (Figure 4H). Interestingly, we found that either hypoxia treatment or overexpression of SIAH2 promoted the interaction between DBC1 and SIAH2, but inhibited the interaction of DBC1 with OTUD5 (Figure 4I, Figure 4-figure supplement 1D). Overall, our results suggest that SIAH2 and OTUD5 competitively interact with DBC1 in response to hypoxia (Figure 4J), cooperatively regulating reversible DBC1 ubiquitination and stability to orchestrate DBC1 function.

**Figure 4.**
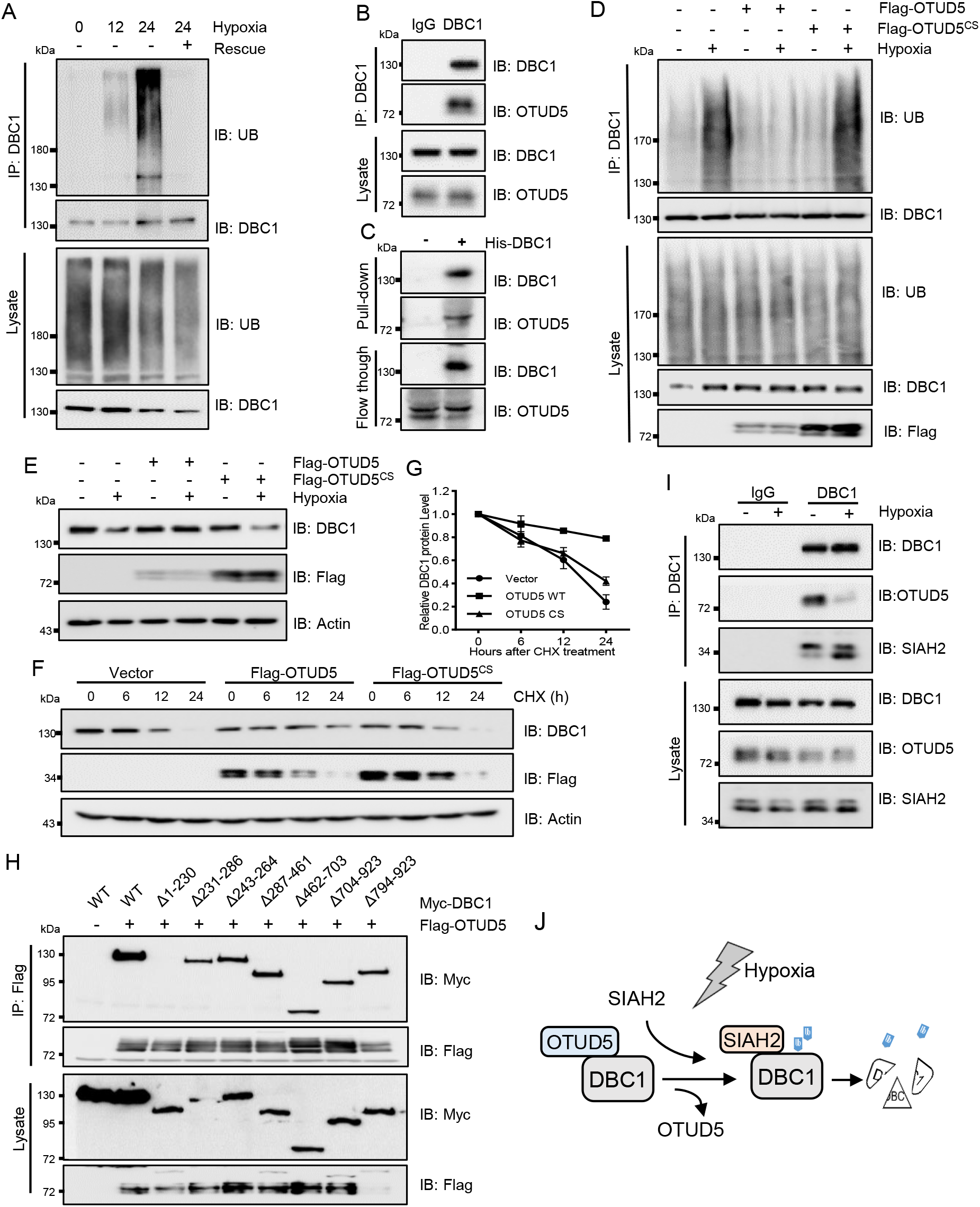
OTUD5 interacts with DBC1 and regulates its stability. **A,** Co-IP analysis showing the ubiquitination level of endogenous DBC1 in MDA-MB-231 cells under hypoxia for 12 h and 24 h and then recovered to normoxia for 6 h. **B,** Co-IP analysis showing the endogenous interaction between OTUD5 and DBC1 in MDA-MB-231 cells. **C,** GST pulldown analysis showing the interaction between His-tagged DBC1 and endogenous OTUD5 immunoprecipitated from MDA-MB-231 cells. **D,** Co-IP analysis showing the ubiquitination level of endogenous DBC1 in HeLa cells transfected with wild-type Falg-OTUD5 or Flag-OTUD5 ^CS^ under hypoxia. **E,** Western blotting analysis showing the protein level of endogenous DBC1 in HeLa cells transfected with wild-type Falg-OTUD5 or Flag-OTUD5 ^CS^ under hypoxia. **F-G,** Western blotting analysis showing the half-life of endogenous DBC1 in HeLa cells transfected with wild-type Flag-OTUD5 or Flag-OTUD5^CS^ and then treated with CHX (10 μM) under hypoxia. F, Quantification of DBC1 protein levels as shown in (**G**). **H,** Mapping the domain of DBC1 interacting with Flag-OTUD5 in HEK293T cells cotransfected with full-length or truncated forms of Myc-DBC1 and Flag--OTUD5. **I,** Co-IP analysis showing the endogenous interaction between DBC1 and SIAH2 or OTUD5 in MDA-MB-231 cells under hypoxia for 24 h. **J,** Schematic model presenting the competitive interaction of SIAH2 and OTUD5 with DBC1 under hypoxia.

### Hypoxia regulates tumor progression via the SIAH2-DBC1 axis

As suggested in Figure 1, hypoxia-induced DBC1 degradation might regulate pathways associated with tumor cell growth, such as SIRT1 and p53 signaling pathways. Therefore, we examined whether SIAH2-mediated DBC1 ubiquitination was responsible for tumor progression. Our results showed that knockout of *Siah2* attenuated the reduction of p53 pathway activity caused by hypoxia (Figure 5A). In addition, deletion of *SIAH2* promoted cell apoptosis under hypoxia, which could be rescued by simultaneous knockout of *DBC1* (Figure 5B, C and Supplementary file 5). The clone formation assay further revealed that cell growth was suppressed when SIAH2 was deleted under hypoxia, whereas double knockout of *Siah2* and *DBC1* almost did not inhibit cell growth (Figure 5D, E and Supplementary file 5). Transwell and scratch assays also revealed that knockout of *Siah2* inhibited cell migration under hypoxia, rather than double knockout of *Siah2* and *DBC1* (Figure 5F-I and Supplementary file 5). These results suggest that SIAH2 promotes cell growth, proliferation and migration in response to hypoxia by ubiquitinating DBC1 to induce DBC1 degradation.

**Figure 5.**
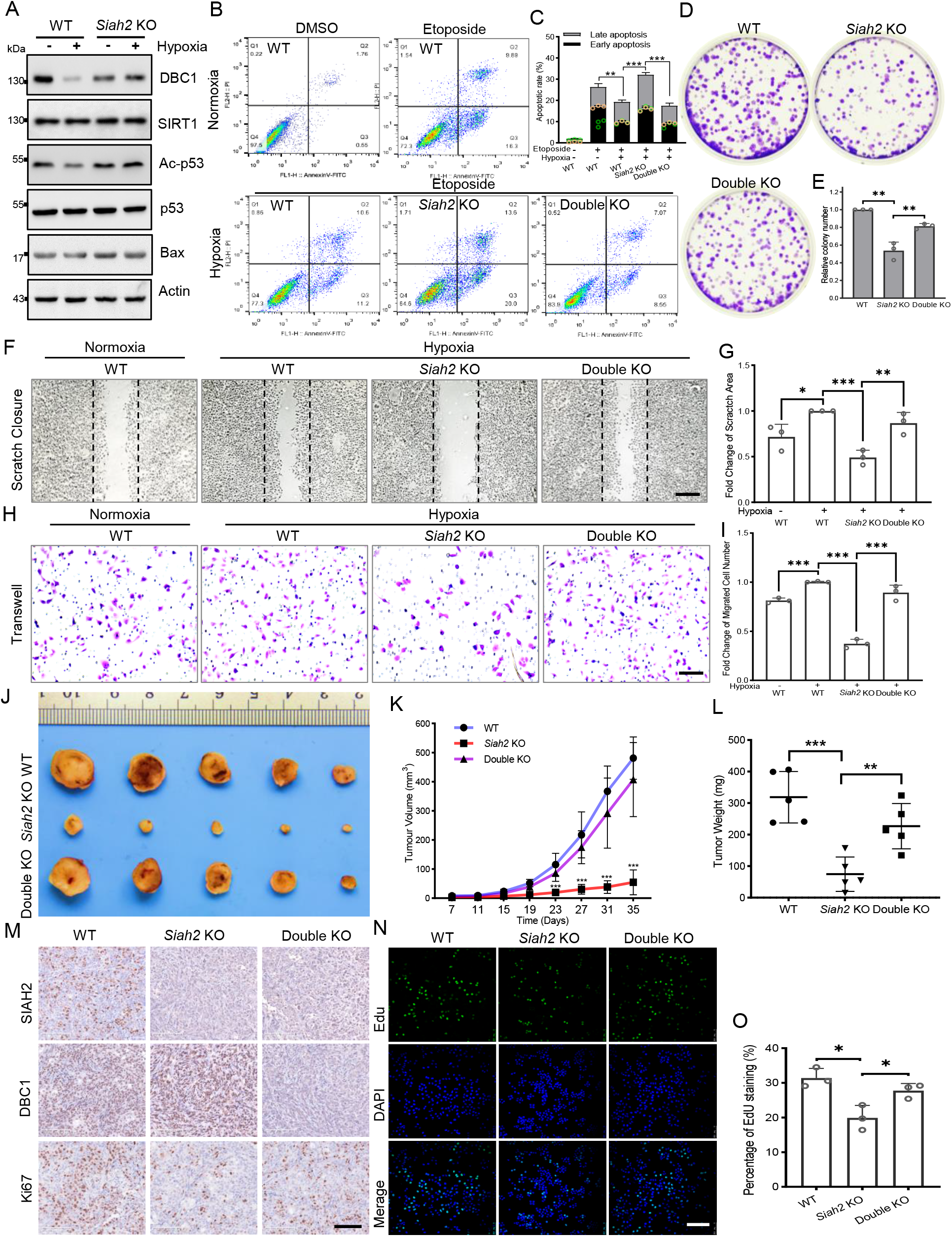
Hypoxia regulates tumor progression via the SIAH2-DBC1 axis. **A,** Western blotting analysis showing the acetylation level of p53 and the protein level of p53 pathway-associated proteins in wild-type and *Siah2* knockout MDA-MB-231 cells. **B,** Apoptosis analysis of wild-type, *Siah2* knockout and *DBC1/Siah2* double knockout MDA-MB-231 cells treated with etoposide (50 μM) under hypoxia. **C,** The apoptosis ratio from (**B**) was quantified (mean ± SEM; n=3 biologically independent samples; *P < 0.05, **P < 0.01). **D,** Colony formation analysis of wild-type, *Siah2* knockout, and *Siah2/DBC1* double knockout MDA-MB-231 cells under hypoxia. **E,** Colony numbers from (**D**) were quantified (mean ± SEM; n=3 biologically independent samples; *P < 0.05, **P < 0.01, ***P < 0.001). **F-G,** Scratching analysis of wild-type, *Siah2* knockout, and *Siah2/DBC1* double knockout MDA-MB-231 cells under hypoxia. scale bars, 200 μm. Data shown as the mean ± SEM.; n=3 biologically independent samples, *P < 0.05, **P < 0.01. **H-I,** Transwell analysis of wild-type, *Siah2* knockout, *and Siah2/DBC1* double knockout MDA-MB-231 cells under hypoxia. scale bars, 100 μm. Data are shown as the mean ± SEM; n=3 biologically independent samples, *P < 0.05, **P < 0.01. **J-L,** Tumor images (**J**), tumor growth curves (**K**) and tumor weight (**L**) from mice subcutaneously injected with *Siah2* knockout or *Siah2/DBC1* double knockout MDA-MB-231 cells. *P < 0.05; **P < 0.01; ***P < 0.001; Two-way ANOVA. **M,** Immunohistochemical analysis of *Siah2* knockout and *Siah2/DBC1* double knockout xenograft tumor tissues with the indicated antibodies. scale bars, 100 μm. **N,** EdU proliferation analysis of different tumor cells isolated from Siah2 knockout or Siah2/DBC1 double knockout xenografts. scale bars, 200 μm. **O,** The percentage of EdU-positive cells from (**N**) was quantified (mean ± SEM; n=3 biologically independent samples; *P < 0.05, **P < 0.01).

To validate the effects of SIAH2-mediated DBC1 stability on tumorigenesis and tumor progression *in vivo*, we implanted *Siah2* knockout and *Siah2/DBC1* double knockout MDA-MB-231 cells into the mammary fat pads of BALB/c nude mice and then monitored tumor growth. The results showed that both tumor volume and tumor weight in the nude mice transplanted with *Siah2* knockout cells were decreased, whereas concurrent deficiency of SIAH2 and DBC1 restored tumor growth to a certain extent (Figure 5J-L and Supplementary file 5). Immunohistochemical analysis revealed that the number of Ki67-positive proliferative cells in *Siah2* knockout-implanted tumors was decreased compared with that in both wild-type and *Siah2/DBC1* double knockout xenografts (Figure 5M). Furthermore, the EdU assays that using cells isolated from the xenografts of animals also showed similar results (Figure 5N, O and Supplementary file 5). Collectively, these results demonstrate the critical role of the SIAH2-DBC1 axis in promoting tumor progression.

### Correlation of SIAH2 and DBC1 expression with tumor progression in breast cancer

To evaluate whether the mechanism by which the SIAH2-DBC1 axis regulates tumor progression is relevant to human tumorigenesis and tumor progression. We analyzed RNA sequencing data in TCGA (Cancer Genome Atlas) using TIMER, and the results demonstrated that the expression level of SIAH2 was significantly higher in most tumors than in adjacent normal tissues (Figure 6A). In particular, the expression of SIAH2 in breast cancers was higher than that in normal samples (Figure 6B). Analyzing the data from TCGA, we found that SIAH2 was positively correlated with tumor stage and number of lymph nodes, indicative of tumor malignancy (Figure 6C, D). To further explore the clinical relevance of SIAH2-mediated DBC1 degradation, we analyzed the expression of SIAH2 and DBC1 in breast cancer tissue microarrays of patients. The IHC results showed that DBC1 and SIAH2 were negatively correlated in this cohort, coincident with our finding that DBC1 was the substrate of SIAH2 (Figure 6E, F). Statistical analyses of IHC suggested that SIAH2 was significantly upregulated in human breast tumor tissues (Figure 6G), which further validated our results from TCGA analyses. Furthermore, the upregulation of SIAH2 was found to be positively correlated with clinical stage and the percentage of the Ki67-positive cell population (Figure 6H, I), and low-level DBC1 expression was detrimental to the survival rate of breast cancer patients (Figure 6J). These data reveal that under pathological conditions, SIAH2-mediated DBC1 ubiquitination and degradation are beneficial for tumor progression.

**Figure 6.**
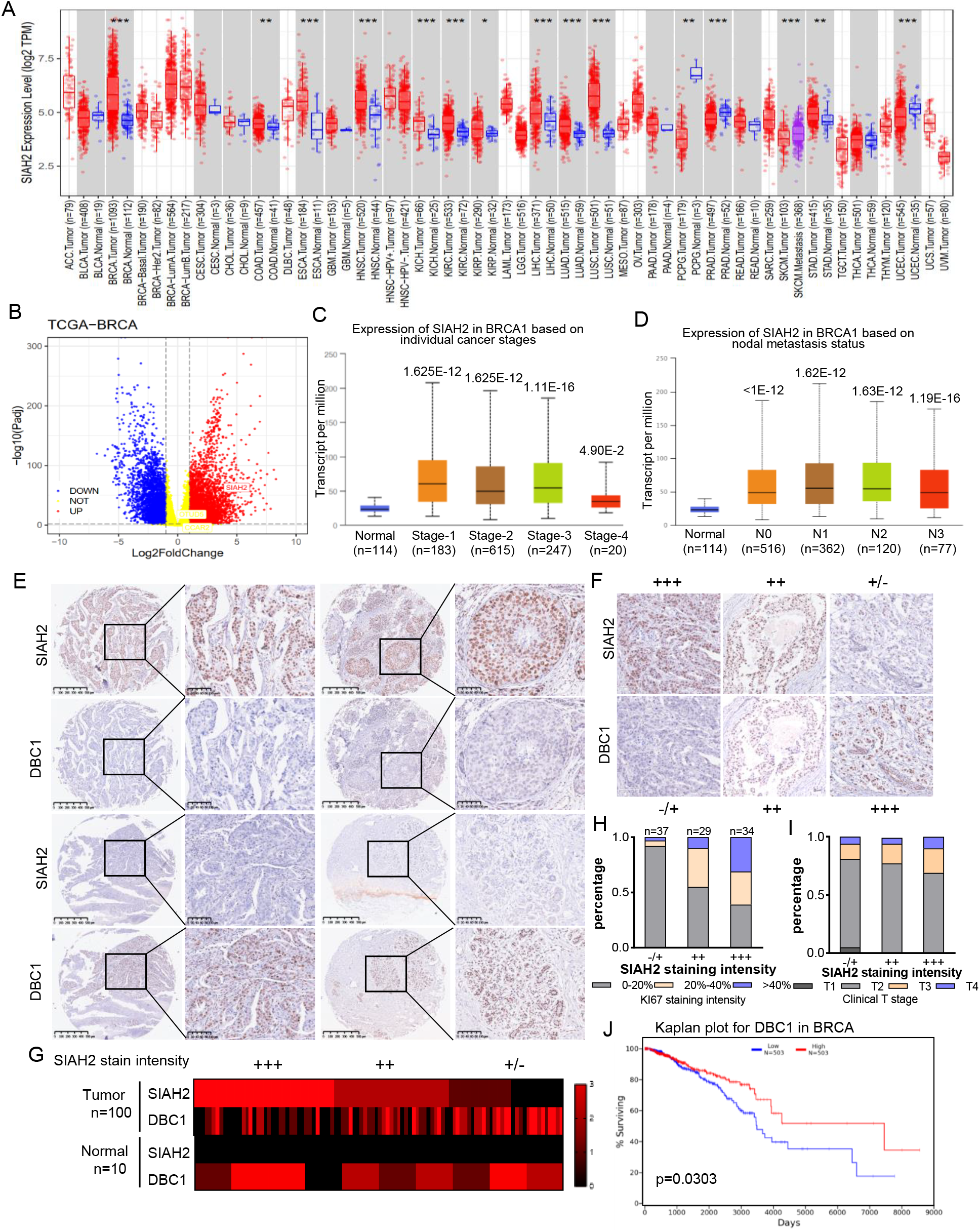
Correlation of SIAH2 and DBC1 expression with tumor progression in BRCA. **A,** SIAH2 expression level in different human cancers from TCGA data in TIMER. *P < 0.05, **P< 0.01, ***P < 0.001. **B,** Volcano plot analysis of SIAH2 expression levels in breast cancer. **C,** Relative expression level of SIAH2 in tumor stage (stage 1, stage 2, stage 3, stage 4 or stage 5) from BRCA patients. **D,** Relative expression level of SIAH2 in nodal metastasis status (normal, N0 or N1) from BRCA patients. **E,** Immunohistochemistry staining analysis of the expression levels of SIAH2 and DBC1 in a series of breast cancer patient tissue microarrays. scale bars, (left) 200 μm; (right) 100 μm.) **F,** Expression levels of SIAH2 in breast cancer tissues were classified into three grades (negative, +, ++, or +++) according to the percentage of immunopositive cells and immunostaining intensity. scale bars, 100 μm. **G,** Heatmap of the expression levels of SIAH2 and DBC1 in human normal breast tissues and breast tumor tissues. **H,** Statistical analysis of correlations between SIAH2 expression level and Ki67-positive stage. **I,** Statistical analysis of correlations between SIAH2 expression level and clinical T stage. **J,** Kaplan-Meier survival curves of breast cancer patients with DBC1 high expression and DBC1 low expression in breast tissues.

**Figure 7.**
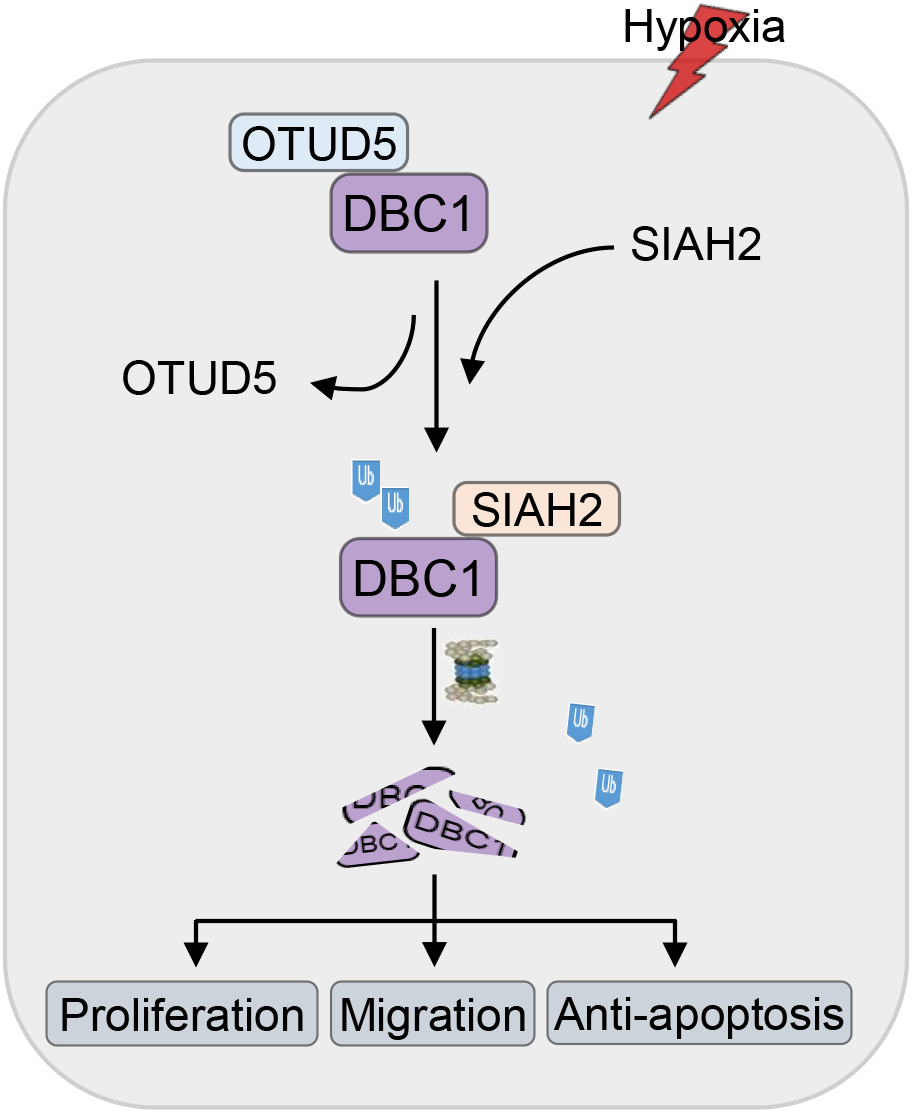
The proposed mechanism by which hypoxia regulates tumor progression through the SIAH2-DBC1 pathway. Under normoxic conditions, the deubiquitinase OTUD5 contacts DBC1 to form a complex. In response to hypoxia, the E3 ubiquitin ligase SIAH2 competitively binds and ubiquitinates DBC1 with OTUD5, resulting in the degradation of DBC1 through the ubiquitin-proteasome system. In the hypoxic microenvironment of the tumor, SIAH2-mediated DBC1 degradation promotes cell migration and proliferation.

## Discussion

It has been documented that DBC1, a specific nuclear protein containing multifunctional domains, participates in the positive and negative regulation of multiple signaling pathways. Several findings have suggested that DBC1 acts as a natural and endogenous inhibitor of SIRT1, and DBC1 deletion increases the SIRT1-p53 interaction and represses p53 transcriptional activity to inhibit apoptosis and promote tumorigenesis (Kim, Chen, and Lou 2008; Zhao et al. 2008; Noh et al. 2013; Qin et al. 2015; Akande et al. 2019). Given the importance of DBC1 function, the regulatory mechanisms by which DBC1 protein stability is regulated remain unclear. Here, we showed that the protein level of DBC1 was degraded under hypoxic stress, which was dynamically regulated by the E3 ubiquitin ligase SIAH2 and deubiquitinase OTUD5. Additionally, knockout of *Siah2* promoted apoptosis and reduced cell migration and tumor proliferation, whereas double knockout of *DBC1* and *Siah2* partially rescued cell proliferation and tumorigenesis. Importantly, DBC1 is negatively correlated with SIAH2 expression levels in human breast tumors, suggesting that the SIAH2–DBC1 axis pathway may play a key role in human breast cancer. These results provide novel insights into the metastatic mechanisms of human breast cancer.

An abundance of evidence has shown that SIAH2 mediates the ubiquitination and degradation of substrates, including PI3K, LATS2, Spry2, ACK1 and TYK2 (Chan et al. 2017; Ma et al. 2016; Buchwald et al. 2013; Nadeau et al. 2007; Muller et al. 2014), in response to hypoxic stress to modulate multiple signaling pathways, such as the Hippo pathway, Ras signaling pathway and STAT3 pathway. In general, DBC1 activity is regulated by multiple posttranslational modifications, including phosphorylation (Magni et al. 2015; Zannini et al. 2012), acetylation (Zheng et al. 2013; Rajendran et al. 2019) and sumolytion (Park et al. 2014). However, it remains unclear whether DBC1 function can be regulated by ubiquitination modification. Our group demonstrated for the first time that SIAH2 can also ubiquitinate DBC1 at Lys287 by binding to the N-terminus of DBC1, resulting in DBC1 degradation by the ubiquitin–proteasome pathway, and that deletion of *SIAH2* will block DBC1 degradation in response to hypoxic stress. Collectively, we identify a novel mechanism by which SIAH2 regulates DBC1 protein stability in response to hypoxic stress, contributing to tumorigenesis and tumor progression.

It has been reported that DBC1 cooperates with SIRT1 to mediate different physiological functions within mammalian cells. For example, stimulation of DBC1 transcription inhibits SIRT1 activity, contributing to TGF-β-induced epithelial-mesenchymal transition (Chen et al. 2021). Additionally, SIRT7 represses DBC1 transcription to promote thyroid tumorigenesis by binding to the promoter of DBC1 (Li et al. 2019). Recently, DBC1 was found to play a role in upregulating glucose homeostasis-related genes, which are implicated in Type 2 diabetes pathogenesis (Basu et al. 2020). Strikingly, our findings reveal that SIAH2-mediated DBC1 ubiquitination under hypoxia regulates cell proliferation and tumorigenesis. We also found that the decrease in p53 acetylation was accompanied by DBC1 degradation; thus, we further validated that the p53 signaling pathway is the downstream mechanism contributing to DBC1-regulated breast tumor progression.

In general, ubiquitination is a dynamic and reversible process cooperatively regulated by E3 ligases and DUBs (Li and Reverter 2021). In this study, we identified that the deubiquitinase OTUD5 could specifically cleave the polyubiquitin chains of DBC1 once hypoxic stress was removed. If OTUD5 was overexpressed under hypoxia, the ubiquitination and degradation of DBC1 would be inhibited. Interestingly, we further confirmed that SIAH2 and OTUD5 competitively bind to DBC1 at the same N-terminal region, and hypoxia promotes the interaction of DBC1 with SIAH2 rather than OTUD5, resulting in ubiquitination and degradation of DBC1 to promote tumor progression. It is well known that OTUD5 controls cell survival and cell proliferation by deubiquitinating substrates (Zhang et al. 2021; Guo et al. 2021; Cho et al. 2021; Li et al. 2020; de Vivo et al. 2019; Park et al. 2015; Luo et al. 2013; Kayagaki et al. 2007). Based on our findings and others’, we speculate that the activation of OTUD5-mediated DBC1 deubiquitination may suppress tumor growth, which may provide a novel target for clinical tumor therapy. Moreover, we still need to further investigate the details of how the OTUD5-DBC1 complex dissociates under hypoxia to modulate DBC1 stability.

Taken together, our study revealed that hypoxia stimulates SIAH2 to ubiquitinate DBC1 and inhibit OTUD5-mediated DBC1 deubiquitination, resulting in DBC1 degradation through the ubiquitin–proteasome pathway. Our results further address the importance of the SIAH2-DBC1 axis in promoting tumor cell survival and migration. In conclusion, we uncovered a complete and detailed dynamic regulatory mechanism by which DBC1 protein ubiquitination and stability are regulated under hypoxic stress. It is of great significance to deeply understand the function of SIAH2 and DBC1 and the role of hypoxia in regulating tumorigenesis and tumor progression, which provides a solid theoretical basis for cancer treatment.

## Acknowledgements

We are grateful to Dr. Wenhui Zhao for providing us with the OTUD5 plasmids at the Center for Peking University Health Science Center. We are grateful to Dr. Changhai Tian from the University of Kentucky College of Medicin for critical reading of the manuscript. This research was supported by grant 2019YFA0508603 from the Ministry of Science and Technology of China and grants 32030026, 91754114 and 31271529 from the Natural Science Foundation of China to Yushan Zhu and Quan Chen.

## Author contributions

QL conceived and designed the experiments with YZ; QL performed most of the experiments and data analysis with assistance from QL, JF, YZ, LL, BM, HC, TZ, JZ, CM, LC, HL, YL, JS, GC, YL and QC; QL, QL and YZ wrote the manuscript. All authors provided intellectual input to the manuscript.

## Materials and Methods

### Construction of plasmids

The expression plasmids for human SIAH2, DBC1, SIRT1 and ubiquitin were generated by amplifying the corresponding cDNA by PCR and cloning it into pcDNA4-TO-Myc-His-B, pCMV-3xFLAG, pEGFP-C1, pRK5-HA, pGEX4T1 or PET28a expression vectors. Site-specific mutants and special domain deletion mutants were generated using TransStart^®^ FastPfu DNA Polymerase (Transgene, Cat# AP221-11) according to the manufacturer’s protocols.

### Cell culture and transfection

HEK293T cells, HeLa cells, MDA-MB-231 cells, MCF7 cells, SH-SY5Y cells, U2OS cells, SW480 cells, HCT116 cells, PC3 cells and HepG2 cells were cultured in GIBCO Dulbecco’s modified Eagle’s medium (DMEM) supplemented with 10% fetal bovine serum (FBS, Lanzhou Bailing) and 100 units/mL penicillin and 100 mg/mL streptomycin in a 5% CO_2_ incubator at 37 °C. The hypoxia gas mixture containing 1% O_2_, 5% CO_2_, and 94% N2 was flushed into a hypoxic chamber (Billups-Rothenberg) to achieve hypoxic culture conditions. For HeLa and HEK293T cells, plasmids were transfected with polyethyleneimine (PEI; Polysciences, Cat# 23966) according to the manufacturer’s protocols. For MDA-MB-231 cells, plasmids were transfected with Lipofectamine 3000 (Invitrogen, Cat# L3000015) according to the manufacturer’s instructions.

### Stable knockdown and knockout cells generation

To generate HeLa and MDA-MB-231 cell lines with stable knockdown of Siah2 and DBC1, oligonucleotides were cloned into pLKO.1, and then the acquired plasmid was cotransfected into HEK293T cells with lentiviral packaging plasmids psPAX2 and pMD2. G for lentivirus production. After infection, MDA-MB-231 cells were selected with 2.5 μg/ml puromycin in culture medium. The single oligonucleotide pair used was as follows: SIAH2 #1 (5’-CCGGGCTGGCTAATAGACACTGAATCTCGAGATTCAGTGTCTATTAGCCAGCTTTTT-3’); SIAH2 #2 (5’-CCGGCCATGATGTGACTTTCGTAAACTCGAGTTTACGAAAGTCACATCATGGTTTTTG-3’); DBC1 #1 (5’-CCGGGCCAAAGGAAAGGATCTCTTTCTCGAGAAAGAGATCCTTTCCTTTGGCTTTTTG-3’); and DBC1 #2 (5’-CCGGGCATTGATTTGAGCGGCTGTACTCGAGTACAGCCGCTCAAATCAATGCTTTTTG-3’). To generate Siah2 and DBC1 knockout HeLa and MDA-MB-231 cell lines, oligonucleotides were cloned into Lenti-CRISPR, and then the acquired plasmid was cotransfected into HEK293T cells with lentiviral packaging plasmids psPAX2 and pMD2. G for lentivirus production. After infection, stable HeLa and MDA-MB-231 cells were selected with 2.5 μg/mL puromycin, and then single clones were picked. The knockout clones were confirmed by sequencing the edited genomic regions after PCR amplification and by western blotting.

### Coimmunoprecipitation and pull-down assay

After the described treatment, cells were collected and lysed in 0.8 mL IP lysis buffer (150 mM NaCl, 20 mM Tris, pH 7.4, 1 mM EGTA, 1 mM EDTA, 1% NP-40, 10% glycerol) containing protease inhibitors (1:100, Roche) for 45 min on a rotor at 4 °C. After centrifugation at 12,000 g for 10 min, the supernatant was immunoprecipitated with 1.5 μg of specific antibodies overnight at 4 °C. Protein A/G agarose beads (15-30 μL, Santa Cruz) were washed and then added for another 2 h. The precipitants were washed seven times with wash buffer, and the immune complexes were boiled with loading buffer and analysed. GST-SIAH2 and His-DBC1 proteins were generated in *E. coli* and incubated *in vitro* overnight. Then, 25 μL GST beads (GE Healthcare) were added to the system for 2 h on a rotor at 4 °C.

### Ubiquitination assay

*In vivo* ubiquitination assays Cells were transiently transfected with plasmids expressing HA-ubiquitin or special site mutations and Myc-DBC1 together with Flag-SIAH2 or Flag-SIAH2^RM^ 24 hours after transfection. Cells were treated with 10 μM MG132 (Selleck, S2619) for 6 h before collection. Cells were washed with cold PBS and then lysed in 200 μl of denaturing buffer (150 mM Tris-HCl, pH 7.4, 1% SDS) by sonication and boiling for 15 min. Lysates were made up to 1 ml with regular lysis buffer and immunoprecipitated with 2 μg anti-c-Myc antibody at 4 °C overnight, washed 3 times with cold lysis buffer and then analysed by SDS–PAGE. For the DBC1 endogenous ubiquitination assay, cells were treated with 10 μM MG132 for 6 h before collection. Lysates were immunoprecipitated using 2 μg anti-UB antibody or anti-DBC1 antibody and subjected to ubiquitination analysis by western blotting with anti-DBC1 or anti-ubiquitin antibody. *In vitro* ubiquitination assays were carried out in ubiquitination buffer (50 mM Tris, pH 7.4, 5 mM MgCl_2_, 2 mM dithiothreitol) with human recombinant E1 (100 ng, Upstate), human recombinant E2 UbcH5c (200 ng, Upstate), and His-tagged ubiquitin (10 μg, Upstate). GST-SIAH2, GST-SIAH2^RM^ and His-DBC1 were expressed and purified from Escherichia coli BL21 cells. Two micrograms of GST, GST-SIAH2 or GST-SIAH2^RM^ protein was used in the corresponding ubiquitination reactions. Reactions (total volume 30 μL) were incubated at 37 °C for 2 h and subjected to ubiquitination analysis by western blotting using anti-DBC1 antibody.

### Flow cytometry

After the described treatment, cells were collected and washed twice with prewarmed FBS-free DMEM and then stained with PI and Annexin V (Thermo Fisher) for 20 min at 37 °C. After staining, cells were washed twice with prewarmed PBS for analysis with a flow cytometer (BD Calibur).

### Western blotting

Cells were collected and washed with PBS and then lysed in 1% SDS lysis buffer or NP-40 lysis buffer (150 mM NaCl, 20 mM Tris, pH 7.4, 1 mM EGTA, 1 mM EDTA, 1% NP-40, 10% glycerol) containing protease inhibitors (1:100, Roche) for 45 min on a rotor at 4 °C. After centrifugation at 12,000 g for 10 min, the supernatant was boiled with loading buffer. Cell lysates containing equivalent protein quantities were subjected to 6% or 10% SDS-PAGE, transferred to nitrocellulose membranes, and then incubated with 5% milk for 2 h at room temperature. Then, the membranes were probed with related primary antibodies at 4 °C, followed by the appropriate HRP-conjugated secondary antibodies (KPL). Immunoreactive bands were checked by a chemiluminescence kit (En-green Biosystem) and visualized by a chemiluminescence imager (JUNYI). The following antibodies were used: Flag-M2 (1:2000, Sigma), Myc (1:1000, Santa Cruz Biotech.), HA (1:1000, Santa Cruz Biotech.), GFP (1:1000, Santa Cruz Biotech.), His (1:1000, Santa Cruz Biotech.), GST (1:1000, Santa Cruz Biotech.), anti-ACTIN (1:10000, Sigma), anti-mono- and polyubiquitinated conjugate monoclonal antibodies (FK2) (1:1000, Enzo Life Sciences), SIAH2 (1:500 Proteintech), OTUD5 (1:1000 Proteintech & CST), DBC1 (1:1000 Proteintech & CST & Abcam), SIRT1 (1:1000 Abcam), p53 (1:1000 Abcam), acetylated-K382-p53 (1:1000 Abcam), HIF1α (1:1000 Proteintech), BAX (1:1000 Proteintech), p21 (1:1000 Proteintech), and GSK3β (1:1000 Proteintech).

### RNA extraction and real-time PCR

RNA samples were extracted with TRIzol reagent (Sigma T9424), reverse transcription PCR was performed with a Reverse Transcriptase kit (Promega A3803), real-time PCR was performed using Powerup SYBR Green PCR master mix (A25743) and a Step-One Plus realtime PCR machine (Applied Biosystems). Human Actin expression was used for normalization.

### Quantification and statistical analysis

For quantitative analyses of western blots, real-time PCR results or flow cytometry data, values were obtained from three independent experiments. The quantitative data are presented as the means ± SEM. Student’s t test was performed to assess whether significant differences existed between groups. Multiple comparisons were performed with one-way analysis of variance (ANOVA) and Tukey’s post-hoc test. P values < 0.05 were considered statistically significant. The significance level is presented as *P < 0.05, **P < 0.01 and ***P < 0.001. “ns” indicates that no significant difference was found. All analyses were performed using Prism 8.0 (GraphPad Software, Inc., La Jolla, CA).

### Tissue microarrays and immunohistochemistry

The breast cancer tissue microarrays were purchased from US Biomax. These tissue microarrays consisted of 100 analysable cases of invasive breast carcinoma and 10 analysable cases of normal breast tissue. For antigen retrieval, the slides were rehydrated and then treated with 10 mM sodium citrate buffer (pH 6.0) heated for 3 min under pressure. The samples were treated with 3% H_2_O_2_ for 15 min to block endogenous peroxidase activity and then blocked with 5% goat serum for 1 h at room temperature. Then, the tissues were incubated with the indicated antibodies at 4 °C overnight, followed by incubation with HRP-conjugated secondary antibody for 1 h at room temperature. Immunoreactive signals were visualized with a DAB Substrate Kit (MaiXin Bio). Protein expression levels in all the samples were scored on a scale of four grades (negative, +, ++, +++) according to the percentage of immunopositive cells and immunostaining intensity. The χ^2^ test was used for analysis of statistical significance. The following antibodies were used for immunohistochemistry: Ki67 (1:200, Abcam, ab16667), SIAH2 (1:40, Novus Biologicals, NB110-88113) and DBC1 (1:100, Proteintech, 22638-1-AP).

### Xenograft tumorigenesis study

All mouse experiments were approved by the Institutional Animal Care and Use Committee at the College of Life Sciences at Nankai University. MDA-MB-231 breast cancer cells (2 × 10^6^ in 100 μL PBS) were injected subcutaneously into the armpit of six-to eight-week-old female BALB/c nude mice. Tumor size was measured every 3–5 days one week after the implantation, and tumor volume was also analysed by using the formula V = 0.5×L×W2 (V: volume, L: length, W: width). The mice were then sacrificed, and the subcutaneous tumors were surgically removed, weighed and photographed. No statistical method was used to predetermine the sample size for each group. The experiments were not randomized.

### RNA-sequencing analysis

RNA was extracted in biological triplicates using the miRNeasy Mini kit (Qiagen) according to the manufacturer’s instructions. RNA degradation and contamination were monitored on 1% agarose gels. RNA purity was checked using a NanoPhotometer^®^ spectrophotometer (IMPLEN, CA, USA). RNA integrity was assessed using the RNA Nano 6000 Assay Kit of the Bioanalyzer 2100 system (Agilent Technologies, CA, USA). RNA quality control was performed using a fragment analyser and standard or high-sensitivity RNA analysis kits (Labgene; DNF-471-0500 or DNF-472-0500). RNA concentrations were measured using the Quanti-iTTM RiboGreen RNA assay Kit (Life Technologies/Thermo Fisher Scientific). A total of 1000 ng of RNA was utilized for library preparation with the NEBNext^®^ UltraTM RNA Library Prep Kit for Illumina^®^ (NEB, USA) following the manufacturer’s recommendations, and index codes were added to attribute sequences to each sample. Poly-A + RNA was sequenced with a HiSeq SBS Kit v4 (Illumina) on an Illumina HiSeq 2500 using protocols defined by the manufacturer.

Raw data (raw reads) in fastq format were first processed through in-house Perl scripts. In this step, clean data (clean reads) were obtained by removing reads containing adapters, reads containing poly-N and low-quality reads from raw data. At the same time, the Q20, Q30 and GC contents of the clean data were calculated. All downstream analyses were based on clean data with high quality. Reference genome and gene model annotation files were downloaded from the genome website directly. The index of the reference genome was built using Hisat2 v2.2.1, and paired-end clean reads were aligned to the reference genome using Hisat2 v2.2.1. We selected Hisat2 as the mapping tool because Hisat2 can generate a database of splice junctions based on the gene model annotation file and thus a better mapping result than other nonsplice mapping tools. The mapped reads of each sample were assembled by StringTie (v2.1.4) (Mihaela Pertea.et al. 2015) in a reference-based approach. StringTie uses a novel network flow algorithm as well as an optional de novo assembly step to assemble and quantitate full-length transcripts representing multiple splice variants for each gene locus. Differential expression analysis of two conditions/groups (two biological replicates per condition) was performed using the DESeq2 R package (1.30.1). DESeq2 provides statistical routines for determining differential expression in digital gene expression data using a model based on the negative binomial distribution. The resulting P values were adjusted using Benjamini and Hochberg’s approach for controlling the false discovery rate. Genes with an adjusted P value of 0.05 and absolute log2FoldChange of 0.5 were set as the threshold for significantly differential expression.

## Supplementary Figure legends

**Figure 1-figure supplement 1.**
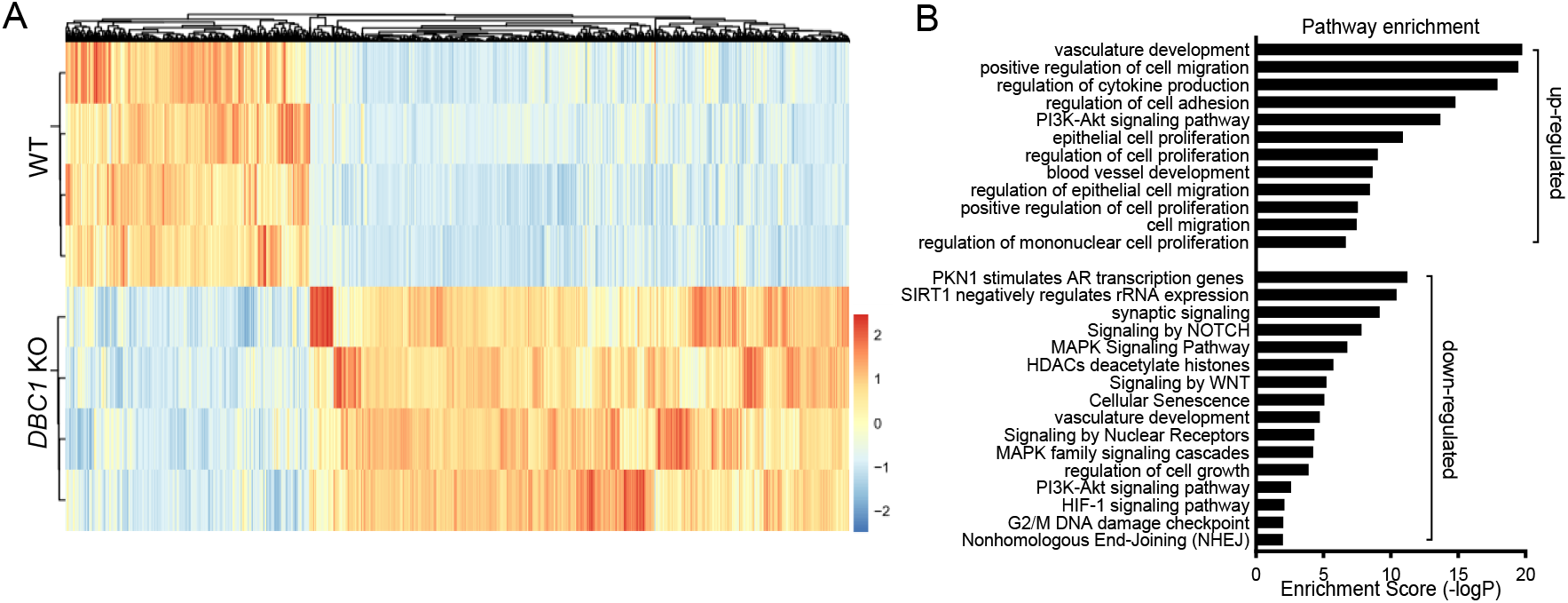
Hypoxia induces DBC1 degradation. **A,** Heatmap of the differentially expressed genes in wild-type and *DBC1* knockout MDA-MB-231 cells. **B,** Pathway enrichment in *DBC1* knockout MDA-MB-231 cells by Metascape.

**Figure 2-figure supplement 1.**
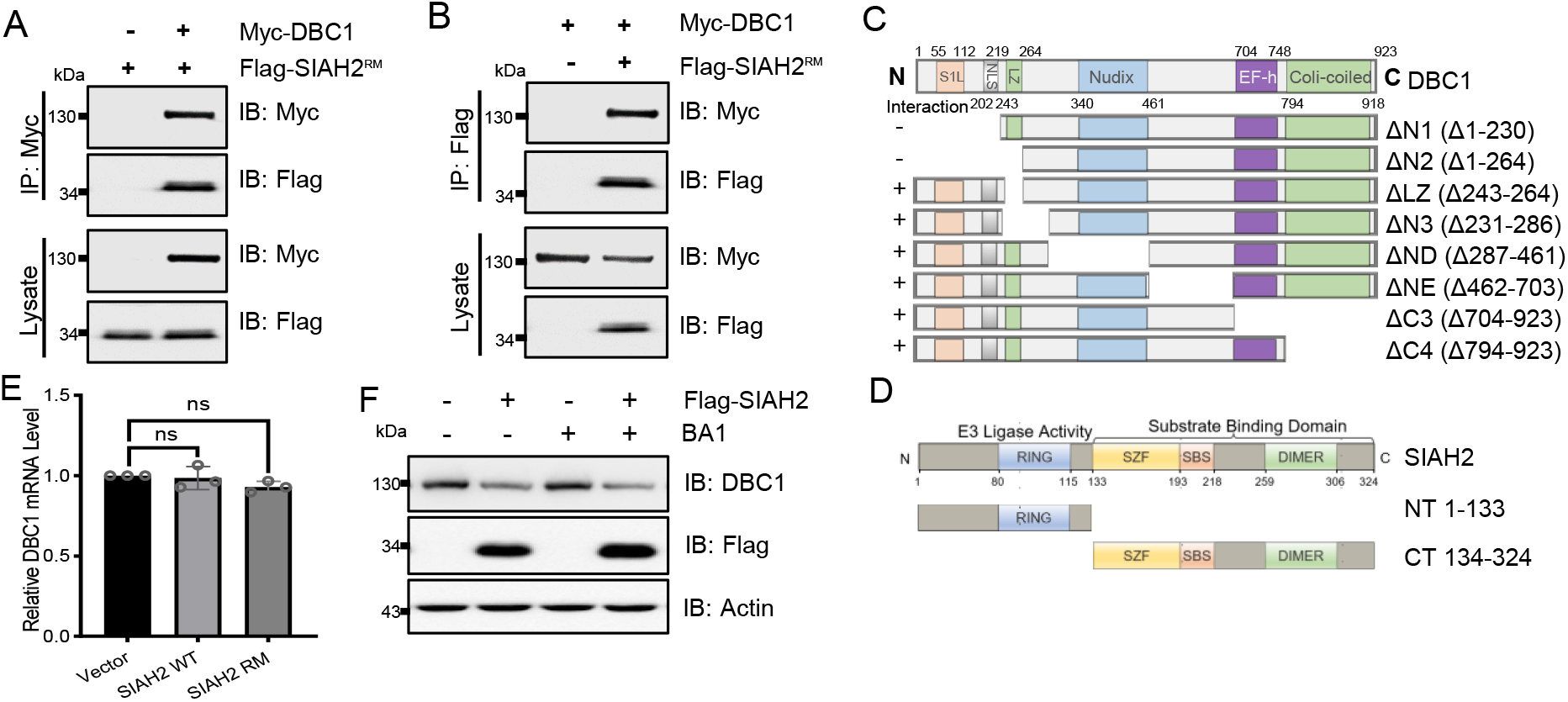
SIAH2 interacts with DBC1 and regulates its stability. **A-B,** Co-IP analysis of the interaction between DBC1 and SIAH2 in HEK293T cells exogenously expressing Myc-DBC1 and Flag-SIAH2^RM^. **C-D,** Models of truncated forms of Myc-DBC1 or Flag-SIAH2. **E,** qRT-PCR analysis of the DBC1 mRNA level in MDA-MB-231 cells. **F,** Western blotting analysis of the DBC1 level in HeLa cells expressing Flag-SIAH2 and treated with the lysosomal inhibitor BA1.

**Figure 3-figure supplement 1.**
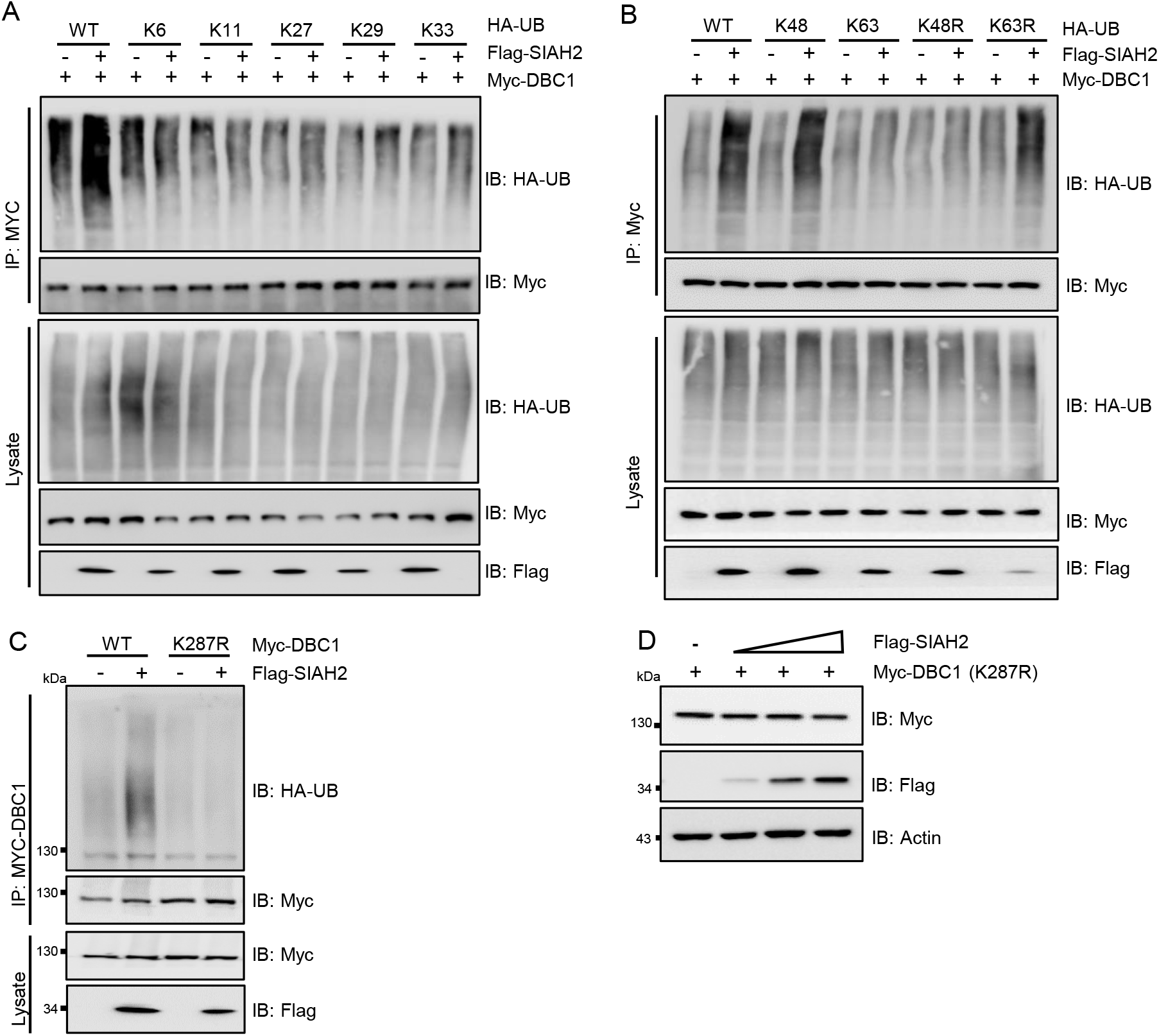
SIAH2 promotes DBC1 polyubiquitination and degradation under hypoxia. **A-B,** Co-IP analysis of the ubiquitination level of DBC1 in HEK293T cells cotransfected with Myc-DBC1, Flag-SIAH2 and wild-type HA-Ub or Ub mutants. **C,** Co-IP analysis of the ubiquitination level of DBC1 in HEK293T cells cotransfected with Flag-SIAH2 and wild-type DBC1 or DBC1-K287R. **D,** Western blotting analysis of the protein level of DBC1-K287R in HeLa cells transfected with SIAH2 at the indicated dosage.

**Figure 4-figure supplement 1.**
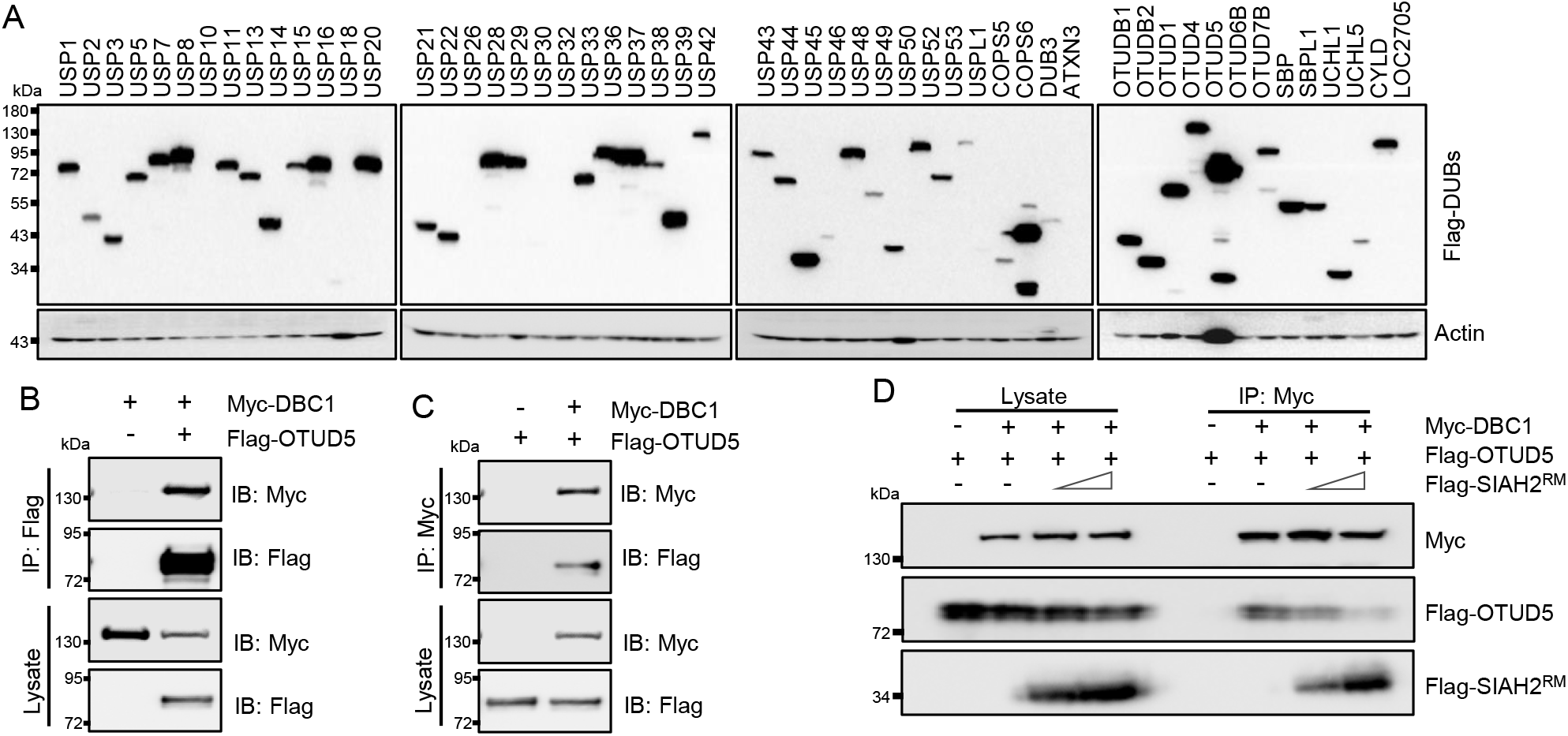
OTUD5 interacts with DBC1 and regulates its stability. **A,** Western blotting analysis of the expression levels of the indicated DUBs in HeLa cells transfected with the indicated Flag-tagged DUBs. **B-C,** Co-IP analysis of the interaction between DBC1 and OTUD5 in HEK293T cells cotransfected with Myc-DBC1 and Flag-OTUD5. **D,** Co-IP analysis of the interaction between DBC1 and SIAH2 and the interaction between DBC1 and OTUD5 in HEK293T cells cotransfected with Myc-DBC1, Flag-OTUD5 and Flag-SIAH2^RM^ at the indicated dosages.

## References

Akande, O. E., P. K. Damle, M. Pop, N. E. Sherman, B. B. Szomju, L. V. Litovchick, and S. R. Grossman. 2019. ‘DBC1 Regulates p53 Stability via Inhibition of CBP-Dependent p53 Polyubiquitination’, Cell Rep, 26: 3323–35 e4.

Basu, S., M. Barad, D. Yadav, A. Nandy, B. Mukherjee, J. Sarkar, P. Chakrabarti, S. Mukhopadhyay, and D. Biswas. 2020. ‘DBC1, p300, HDAC3, and Siah1 coordinately regulate ELL stability and function for expression of its target genes’, Proc Natl Acad Sci U S A, 117: 6509–20.

Buchwald, M., K. Pietschmann, P. Brand, A. Gunther, N. P. Mahajan, T. Heinzel, and O. H. Kramer. 2013. ‘SIAH ubiquitin ligases target the nonreceptor tyrosine kinase ACK1 for ubiquitinylation and proteasomal degradation’, Oncogene, 32: 4913–20.

Chan, L. W., F. Wang, F. Meng, L. Wang, S. C. Wong, J. S. Au, S. Yang, and W. C. Cho. 2017. MiR-30 Family Potentially Targeting PI3K-SIAH2 Predicted Interaction Network Represents a Novel Putative Theranostic Panel in Non-small Cell Lung Cancer’, Front Genet, 8: 8.

Chen, B., W. Dong, T. Shao, X. Miao, Y. Guo, X. Liu, and Y. Feng. 2021. ‘A KDM4-DBC1-SIRT1 Axis Contributes to TGF-b Induced Mesenchymal Transition of Intestinal Epithelial Cells’, Front Cell Dev Biol, 9: 697614.

Cheng, Y. Q., S. B. Wang, J. H. Liu, L. Jin, Y. Liu, C. Y. Li, Y. R. Su, Y. R. Liu, X. Sang, Q. Wan, C. Liu, L. Yang, and Z. C. Wang. 2020. ‘Modifying the tumour microenvironment and reverting tumour cells: New strategies for treating malignant tumours’, Cell Prolif 53: e12865.

Chini, C. C., C. Escande, V. Nin, and E. N. Chini. 2010. ‘HDAC3 is negatively regulated by the nuclear protein DBC1’, J Biol Chem, 285: 40830–7.

Cho, J. H., K. Kim, S. A. Kim, S. Park, B. O. Park, J. H. Kim, S. Y. Kim, M. J. Kwon, M. H. Han, S. B. Lee, B. C. Park, S. G. Park, J. H. Kim, and S. Kim. 2021. ‘Deubiquitinase OTUD5 is a positive regulator of mTORC1 and mTORC2 signaling pathways’, Cell Death Differ, 28: 900–14.

de Vivo, A., A. Sanchez, J. Yegres, J. Kim, S. Emly, and Y. Kee. 2019. ‘The OTUD5-UBR5 complex regulates FACT-mediated transcription at damaged chromatin’, Nucleic Acids Res, 47: 729–46.

Guo, Y., F. Jiang, L. Kong, H. Wu, H. Zhang, X. Chen, J. Zhao, B. Cai, Y. Li, C. Ma, F. Yi, L. Zhang, B. Liu, Y. Zheng, L. Zhang, and C. Gao. 2021. ‘OTUD5 promotes innate antiviral and antitumor immunity through deubiquitinating and stabilizing STING’, Cell Mol Immunol, 18: 1945–55.

Kayagaki, N., Q. Phung, S. Chan, R. Chaudhari, C. Quan, K. M. O’Rourke, M. Eby, E. Pietras, G. Cheng, J. F. Bazan, Z. Zhang, D. Arnott, and V. M. Dixit. 2007. ‘DUBA: a deubiquitinase that regulates type I interferon production’, Science, 318: 1628–32.

Kim, J. E., J. Chen, and Z. Lou. 2008. ‘DBC1 is a negative regulator of SIRT1’, Nature, 451: 583–6.

Lee, J. W., J. Ko, C. Ju, and H. K. Eltzschig. 2019. ‘Hypoxia signaling in human diseases and therapeutic targets’, Exp Mol Med, 51: 1–13.

Li, F., Q. Sun, K. Liu, L. Zhang, N. Lin, K. You, M. Liu, N. Kon, F. Tian, Z. Mao, T. Li, T. Tong, J. Qin, W. Gu, D. Li, and W. Zhao. 2020. ‘OTUD5 cooperates with TRIM25 in transcriptional regulation and tumor progression via deubiquitination activity’, Nat Commun, 11: 4184.

Li, H., Z. Tian, Y. Qu, Q. Yang, H. Guan, B. Shi, M. Ji, and P. Hou. 2019. ‘SIRT7 promotes thyroid tumorigenesis through phosphorylation and activation of Akt and p70S6K1 via DBC1/SIRT1 axis’, Oncogene, 38: 345–59.

Li, Y., and D. Reverter. 2021. ‘Molecular Mechanisms of DUBs Regulation in Signaling and Disease’, Int J Mol Sci, 22.

Luo, J., Z. Lu, X. Lu, L. Chen, J. Cao, S. Zhang, Y. Ling, and X. Zhou. 2013. ‘OTUD5 regulates p53 stability by deubiquitinating p53’, PLoS One, 8: e77682.

Ma, B., Y. Chen, L. Chen, H. Cheng, C. Mu, J. Li, R. Gao, C. Zhou, L. Cao, J. Liu, Y. Zhu, Q. Chen, and S. Wu. 2015. ‘Hypoxia regulates Hippo signalling through the SIAH2 ubiquitin E3 ligase’, Nat Cell Biol, 17: 95–103.

Ma, B., H. Cheng, R. Gao, C. Mu, L. Chen, S. Wu, Q. Chen, and Y. Zhu. 2016. ‘Zyxin-Siah2-Lats2 axis mediates cooperation between Hippo and TGF-beta signalling pathways’, Nat Commun, 7: 11123.

Ma, B., H. Cheng, C. Mu, G. Geng, T. Zhao, Q. Luo, K. Ma, R. Chang, Q. Liu, R. Gao, J. Nie, J. Xie, J. Han, L. Chen, G. Ma, Y. Zhu, and Q. Chen. 2019. ‘The SIAH2-NRF1 axis spatially regulates tumor microenvironment remodeling for tumor progression’, Nat Commun, 10: 1034.

Magni, M., V. Ruscica, M. Restelli, E. Fontanella, G. Buscemi, and L. Zannini. 2015. ‘CCAR2/DBC1 is required for Chk2-dependent KAP1 phosphorylation and repair of DNA damage’, Oncotarget, 6: 17817–31.

Muller, S., Y. Chen, T. Ginter, C. Schafer, M. Buchwald, L. M. Schmitz, J. Klitzsch, A. Schutz, A. Haitel, K. Schmid, R. Moriggl, L. Kenner, K. Friedrich, C. Haan, I. Petersen, T. Heinzel, and O. H. Kramer. 2014. ‘SIAH2 antagonizes TYK2-STAT3 signaling in lung carcinoma cells’, Oncotarget, 5: 3184–96.

Nadeau, R. J., J. L. Toher, X. Yang, D. Kovalenko, and R. Friesel. 2007. ‘Regulation of Sprouty2 stability by mammalian Seven-in-Absentia homolog 2’, J Cell Biochem, 100: 151–60.

Noguchi, A., K. Kikuchi, H. Zheng, H. Takahashi, Y. Miyagi, I. Aoki, and Y. Takano. 2014. ‘SIRT1 expression is associated with a poor prognosis, whereas DBC1 is associated with favorable outcomes in gastric cancer’, Cancer Med, 3: 1553–61.

Noh, S. J., M. J. Kang, K. M. Kim, J. S. Bae, H. S. Park, W. S. Moon, M. J. Chung, H. Lee, D. G. Lee, and K. Y. Jang. 2013. ‘Acetylation status of P53 and the expression of DBC1, SIRT1, and androgen receptor are associated with survival in clear cell renal cell carcinoma patients’, Pathology, 45: 574–80.

Park, J. H., S. W. Lee, S. W. Yang, H. M. Yoo, J. M. Park, M. W. Seong, S. H. Ka, K. H. Oh, Y. J. Jeon, and C. H. Chung. 2014. ‘Modification of DBC1 by SUMO2/3 is crucial for p53-mediated apoptosis in response to DNA damage’, Nat Commun, 5: 5483.

Park, S. Y., H. K. Choi, Y. Choi, S. Kwak, K. C. Choi, and H. G. Yoon. 2015. ‘Deubiquitinase OTUD5 mediates the sequential activation of PDCD5 and p53 in response to genotoxic stress’, Cancer Lett, 357: 419–27.

Qin, B., K. Minter-Dykhouse, J. Yu, J. Zhang, T. Liu, H. Zhang, S. Lee, J. Kim, L. Wang, and Z. Lou. 2015. ‘DBC1 functions as a tumor suppressor by regulating p53 stability’, Cell Rep, 10: 1324–34.

Rajendran, P., G. Johnson, L. Li, Y. S. Chen, M. Dashwood, N. Nguyen, A. Ulusan, F. Ertem, M. Zhang, J. Li, D. Sun, Y. Huang, S. Wang, H. C. Leung, D. Lieberman, L. Beaver, E. Ho, M. Bedford, K. Chang, E. Vilar, and R. Dashwood. 2019. ‘Acetylation of CCAR2 Establishes a BET/BRD9 Acetyl Switch in Response to Combined Deacetylase and Bromodomain Inhibition’, Cancer Res, 79: 918–27.

Singleton, D. C., A. Macann, and W. R. Wilson. 2021. ‘Therapeutic targeting of the hypoxic tumour microenvironment’, Nat Rev Clin Oncol, 18: 751–72.

Won, K. Y., H. Cho, G. Y. Kim, S. J. Lim, G. E. Bae, J. U. Lim, J. Y. Sung, Y. K. Park, Y. W. Kim, and J. Lee. 2015. ‘High DBC1 (CCAR2) expression in gallbladder carcinoma is associated with favorable clinicopathological factors’, Int J Clin Exp Pathol, 8: 11440–5.

Yu, X. M., Y. Liu, T. Jin, J. Liu, J. Wang, C. Ma, and X. L. Pan. 2013. ‘The Expression of SIRT1 and DBC1 in Laryngeal and Hypopharyngeal Carcinomas’, PLoS One, 8: e66975.

Zannini, L., G. Buscemi, J. E. Kim, E. Fontanella, and D. Delia. 2012. ‘DBC1 phosphorylation by ATM/ATR inhibits SIRT1 deacetylase in response to DNA damage’, J Mol Cell Biol, 4: 294–303.

Zhang, Y., Y. Fan, X. Jing, L. Zhao, T. Liu, L. Wang, L. Zhang, S. Gu, X. Zhao, and Y. Teng. 2021. ‘OTUD5-mediated deubiquitination of YAP in macrophage promotes M2 phenotype polarization and favors triple-negative breast cancer progression’, Cancer Lett, 504: 104–15.

Zhao, W., J. P. Kruse, Y. Tang, S. Y. Jung, J. Qin, and W. Gu. 2008. ‘Negative regulation of the deacetylase SIRT1 by DBC1’, Nature, 451: 587–90.

Zheng, H., L. Yang, L. Peng, V. Izumi, J. Koomen, E. Seto, and J. Chen. 2013. ‘hMOF acetylation of DBC1/CCAR2 prevents binding and inhibition of SirT1’, Mol Cell Biol, 33: 4960–70.

